# Development and validation of NAD^+^ metabolism-related gene signature in prognosis prediction of hepatocellular carcinoma

**DOI:** 10.1101/2023.06.25.546472

**Authors:** Yiheng Du

## Abstract

Liver cancer is a prevalent and deadly malignancy worldwide, with a rising incidence rate. In this study, we focused on the role of NAD^+^ metabolism-related genes in liver cancer prognosis prediction. We identified key genes among NAD^+^ metabolism-related genes and explored their relationship with cancer staging and prognosis using gene expression data. A risk score model was constructed to assess cancer risk and survival status. The model demonstrated significant predictive potential for survival outcomes. Furthermore, we analyzed the risk scores in different populations and performed functional enrichment analyses to gain insights into the biological processes involved. Our study highlights the clinical value and significance of NAD^+^ metabolism-related genes in liver cancer. The findings provide a foundation for personalized treatment strategies and open new avenues for further research in liver cancer management.

## Introduction

Liver cancer is considered one of the most common cancers worldwide. It is the only cancer among the top five causes of mortality that exhibits an increasing incidence rate each year (Siegel, Miller and Jemal, 2019). And liver cancer has a more incidence rate in the developing countries (Starley, Calcagno and Harrison, 2010). Risk factors encompass the presence of hepatitis B virus, hepatitis C virus, fatty liver disease, cirrhosis caused by excessive alcohol consumption, smoking, obesity, diabetes, iron overload, and exposure to diverse dietary elements (Center and Jemal, 2011).

In liver cancer research, several potential therapeutic approaches have been discovered, including necroptosis (Gong et al., 2019), pyroptosis (Fang et al., 2020; Yu et al., 2019), autophagy (Degenhardt et al., 2006; Thorburn et al., 2009; Peng et al., 2013), and metastasis (Su et al., 2015; Kenific, Thorburn and Debnath, 2010). These methods hold promise for the treatment of liver cancer. Nicotinamide adenine dinucleotide (NAD^+^) plays a critical role as a mediator in energy metabolism and signal transduction pathways. According to studies (Lv et al., 2020), NAD^+^ metabolism has been shown to drive tumor immune evasion through a CD8^+^ T cell-dependent mechanism (Cairns, Harris and Mak, 2011; Chiarugi et al., 2012). Furthermore, Nicotinamide phosphoribosyltransferase (NAMPT) and NAD^+^ biosynthesis is frequently upregulated in various human malignancies and play a crucial role in tumor initiation, progression, and recurrence (Gujar et al., 2016; Lucena-Cacace et al., 2018; Nacarelli et al., 2019). Therefore, investigating NAD^+^ metabolism-related genes hold the potential to uncover a novel therapeutic approach for liver cancer.

In our study, we performed screening of key genes among NAD^+^ metabolism-related genes and explored the relationship between gene expression levels and cancer staging as well as prognosis in liver cancer. We constructed a risk score to assess cancer risk and survival status. Additionally, we analyzed the risk scores in different populations. Our predictive model demonstrates significant potential in predicting survival outcomes, highlighting its clinical value and significance.

## Method

### Microarray data

In this study, gene expression levels and clinical characteristics of liver cancer patients were obtained from the TCGA database (https://cancergenome.nih.gov/). A total of 424 samples were used, including 374 liver cancer samples and 50 non-tumor tissues. To obtain genes related to NAD^+^-metabolism, 8 gene sets related to NAD^+^-metabolism were obtained from the Explore the Molecular Signatures Database (MSigDB) and subjected to enrichment analysis to identify NAD^+^-metabolism related genes with differential expression for further analysis. These genes were then subjected to interactome analysis to construct a gene interaction network.

### Construction of NAD^+^ metabolism-related prognostic signature for liver cancer

Univariate Cox regression analysis was used to identify NAD^+^ metabolism-related genes that were significantly associated with survival. These genes were then subjected to multivariate Cox regression analysis to identify the optimal NAD^+^ metabolism-related prognostic genes. The TCGA liver cancer dataset was randomly divided into two groups, a training set and a test set. The training set was analyzed using least absolute shrinkage and selection operator (LASSO) Cox regression and machine learning to determine a prognosis-related CARMRs-based predictive signature. A 5-gene signature was ultimately selected based on the regression and the prognostic potential. Finally, a risk score was calculated using the following formula: Risk score = (exprgene1 × Coefgene1) + (exprgene2 × Coefgene2) + … + (exprgenen × Coefgenen), where exprgene is the expression level of the gene and Coefgene is the coefficient for that gene obtained from the regression analysis.

To validate the model, patients were divided into high and low risk groups based on their median risk score. Progress-free survival (PFS) was then analyzed for all samples, the training set, and the test set to evaluate the accuracy of the model, and the relationship between patient survival status and survival time was analyzed with the scoring model. The diagnostic value of the 5 genes was evaluated by plotting survival curves and using ROC curve analysis, and the area under the curve was calculated. The C-index curve was used to estimate the precision of the model. The superiority of the model was demonstrated by performing PCA and t-SNE analysis to distinguish between high and low risk patients at the level of the overall gene, NAD^+^ metabolism-related genes, and the 5 genes selected by the model.

### Independence analysis of clinical traits

To validate the independence of the risk score prediction, a forest plot was used to represent the effectiveness of the risk score and clinical characteristics of different groups in predicting patient survival status. The overall survival was analyzed in patients classified by age (whether over 65 years old), gender, grade, and stage, to determine the effectiveness of the model in predicting different categories of patients.

### Construction of nomogram

We used clinical characteristics (including age, gender, grade, and stage) and risk scores to create a Kaplan-Meier plot to predict the 1-year, 3-year, and 5-year survival of liver cancer patients. In addition, to evaluate the consistency between predicted survival and actual survival, a calibration curve was plotted.

### Evaluation of the 5-gene signature

To verify the contribution of these 5 genes to risk scoring, the expression levels of these 5 genes in high and low risk groups were analyzed, and the expression of these 5 genes in liver cancer patients was analyzed separately.

### Enrichment functional analysis

First, the expression levels of differential genes containing NAD^+^ metabolism-related differences between the high-risk group and the low-risk group were determined, with a filter set at p < 0.05. Then, GO functional enrichment analysis and KEGG (Kanehisa and Goto, 2000) pathway analysis was performed on the expression of differential genes to determine the role of the metabolic pathways controlled by differential genes in liver cancer.

### Immune-related functional analysis

To explore the relationship between risk score and immune cell infiltration, we used algorithms such as TIMER to calculate the abundance of immune cells in the two risk groups. In addition, the immune-related function of the two risk groups was evaluated using the ssGSEA algorithm. Finally, the relationship between risk score and immune-related gene expression was studied to investigate potentially relevant immune-related genes.

### Tumor mutation load and drug sensitivity analysis

The tumor mutational burden (TMB) (Chan, Wolchok and Snyder, 2015) was calculated as the number of mutated bases per million bases for each tumor, based on the somatic mutation data. The maftools package was used to summarize and visualize the mutation data and to evaluate the relationship between risk score and TMB. The Tumor Immune Dysfunction and Exclusion (TIDE) algorithm was used to predict immune response. Then, the oncoPredict package (Maeser, Gruener and Huang, 2021) in R was used to evaluate the sensitivity to chemotherapy drugs, with a filter set at p < 0.001, and the top five drugs with significant correlations were selected for analysis.

### Statistical analysis

GSEA software was used for enrichment analysis to screen for NAD^+^ metabolism-related genes. The gene interaction network was beautified using Cytoscape. Other analyses and plots were all analyzed using R software.

## Result

### Construction and verification of the NAD^+^ metabolism-related genes predictive signature

Using GSEA enrichment analysis, we identified a set of 97 NAD^+^ metabolism-related genes enriched in liver cancer cells. The network of these genes is complex (Figure 1a). To develop predictive features for NAD^+^ metabolism-related genes associated with survival time in liver cancer patients, we randomly classified TCGA-LIHC into training and testing sets. For the training set, we identified 16 NAD^+^ metabolism-related genes with predictive importance through single-variable Cox regression evaluation (Figure 1b). Through Lasso Cox analysis and multivariate Cox analysis, we further identified 5 NAD^+^ metabolism-related genes (Figure 1c, d). These genes show a complex network of relationships with other NAD^+^ metabolism-related genes, suggesting their potential importance in various biological processes. The risk assessment model is as follows: risk score = (0.2753×TXN expression) + (0.2582×PPARG expression) + (0.2364×SLC2A1 expression) +(0.4175×DLAT expression) + (−0.1177×ADH4 expression).

**Figure 1.**
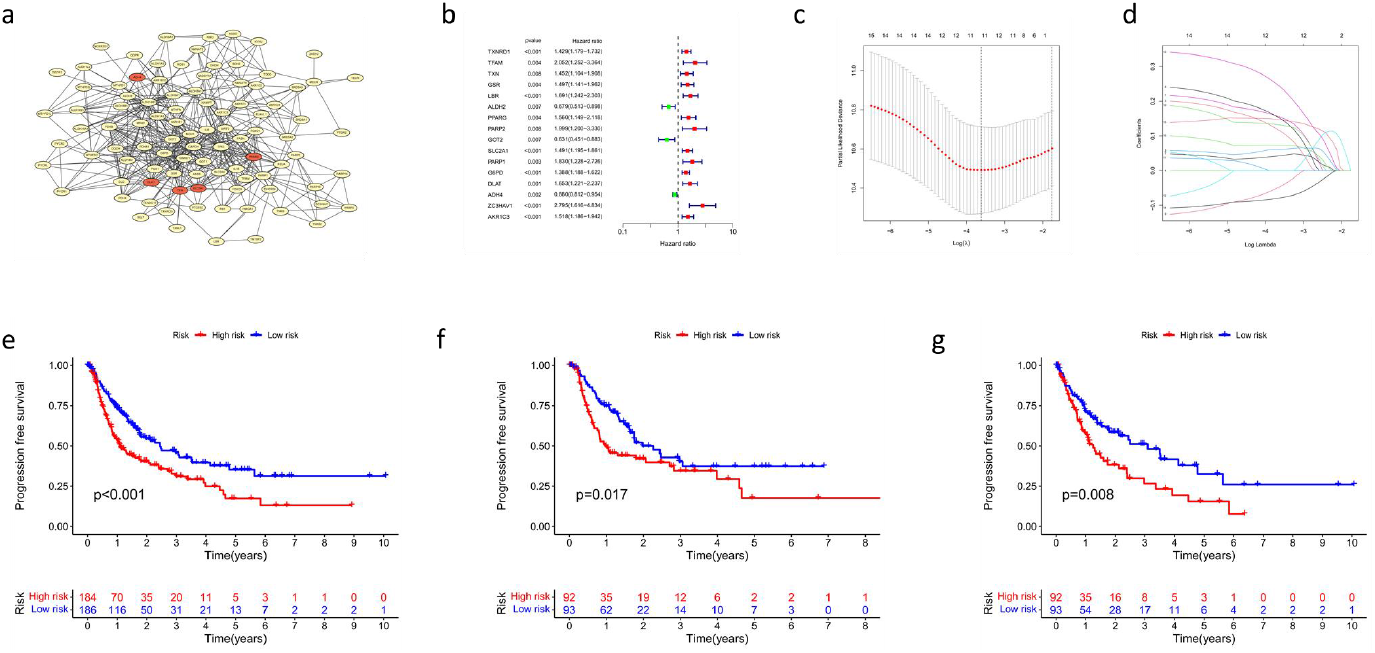
(a) Interacting network of 97 genes related to NAD^+^ metabolism, with the 5 key genes obtained from subsequent screening highlighted in red. (b) 16 NAD metabolism-related genes with strong correlations determined by univariate Cox regression analysis. Genes depicted in red indicate positive associations, while genes in green indicate negative associations. (c) Plot of the multiplication parameters of the Lasso regression model. (d) Relationship between parameters and λ values. (e) Progression-free survival in whole samples. (f) Progression-free survival analysis in the testing set. (g) Progression-free survival in the training set.

Through progression-free survival analysis, it was shown that the risk score model constructed demonstrated significantly lower progression-free survival in the high-risk group compared to the low-risk group in all samples (Figure 1e), testing set (Figure 1f) and training set (Figure 1g). The risk scores and survival status of each patient in all patients, the test set, and the train set were plotted, and it was found that in all three groups, the survival time decreased as the risk score increased (Figure 2a-f).

**Figure 2.**
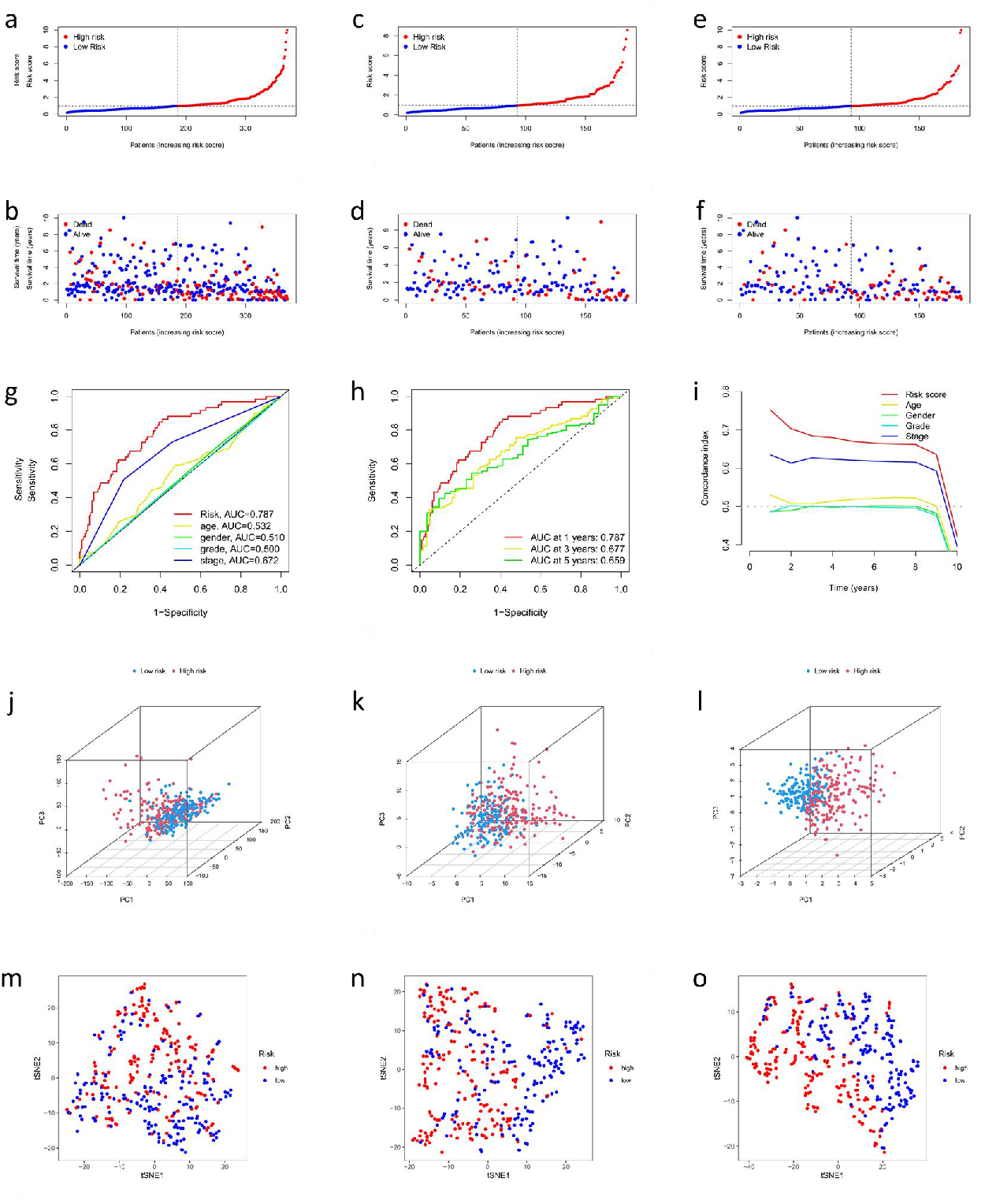
(a) Risk scores of each patient in all patients. (b) The survival time and survival status of each patient in all patients. (c) Risk scores of each patient in the testing set. (d) The survival time and survival status of each patient in the testing set. (e) Risk scores of each patient in the training set. (f) The survival time and survival status of each patient in the training set. (g) ROC curves of risk score, aga, gender, grade, and stage for clinical features in liver cancer patients. (h) Time dependent ROC curves of overall survival at 1, 2- and 3-years. (i) The C-index curve analyzes the consistency index of risk score. (j) PCA analysis differentiating patients with high and low risk scores based on the overall gene expression levels. (k) PCA analysis differentiating patients with high and low risk scores based on the expression levels of NAD metabolism genes. (l) PCA analysis differentiating patients with high and low risk scores based on the expression levels of the five selected NAD metabolism genes. (m) The t-SNE analysis differentiating patients with high and low risk scores based on the overall gene expression levels. (n) The t-SNE analysis differentiating patients with high and low risk scores based on the expression levels of NAD metabolism genes. (o) The t-SNE analysis differentiating patients with high and low risk scores based on the expression levels of the five selected NAD metabolism genes.

To study the prediction accuracy of these 5 gene features, we conducted time-dependent ROC analysis. Additionally, Figure 2g showed the risk score was found to be a better predictor of patient survival than other clinical features such as age, gender, grade, and stage. The AUC of the total survival rate calculated by time (Figure 2h) showed that the gene expression had good prognostic prediction function, with 1 year having an AUC of 0.7, 2 years having an AUC of 0.6, and 3 years having an AUC of 0.6. Next, we constructed the C-index curve (Figure 2i) for risk scores and clinical features, and the results showed that the consistency of risk scores was significantly higher than that of clinical features. Through PCA (Figure 2j-l) and t-SNE (Figure 2m-o) analysis, we found that five genes were able to better distinguish between high-risk and low-risk patients compared to the overall gene and NAD^**+**^ metabolism levels, demonstrating the superiority of the model.

### Independent prognostic analysis

In order to determine whether the risk score could be used as an independent prognostic factor for liver cancer patients, both univariate and multivariate Cox regression analyses were performed. The univariate Cox regression analysis revealed that tumor stage was significantly associated with overall survival time (HR = 1.674, 95% CI: 1.361-2.059, p < 0.001). Furthermore, the risk score was also found to be significantly associated with overall survival time (HR = 1.479, 95% CI: 1.338-1.636, p < 0.001), as indicated in Figure 3a. These findings suggest that both tumor stage and the risk score are independent prognostic factors for liver cancer patients.

**Figure 3.**
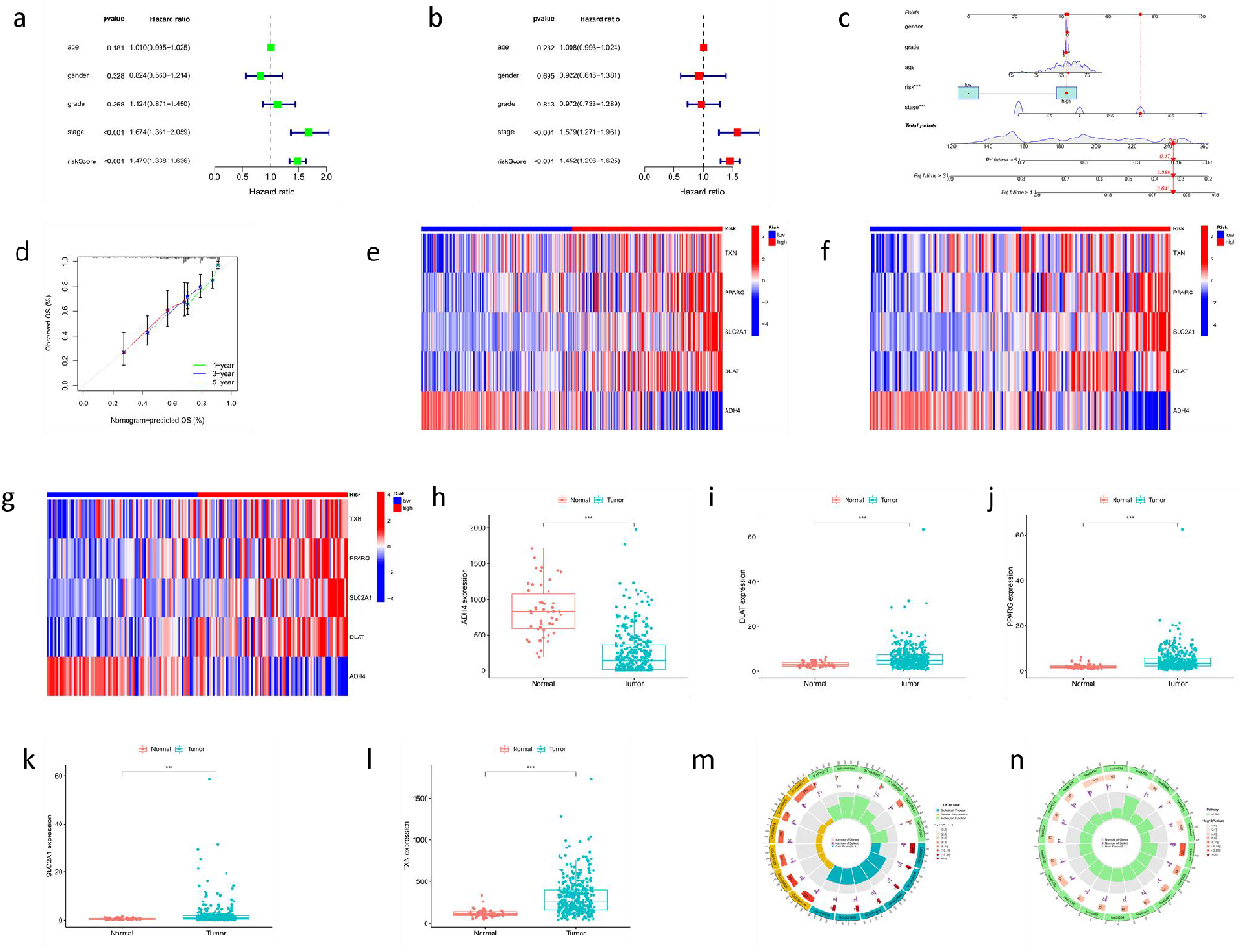
(a) Univariate analysis results. (b) Multivariate analysis results. (c) The nomogram model incorporates the risk factor to predict the survival rates of patients. (d) The calibration curve of the nomogram. (e-g) The relationship between gene expression levels of these five genes and risk score was evaluated in all patient sets, the testing set, and the training set. (h-l) The single gene expression analysis between normal and tumor of the 5 genes (ADH4, DLAT, PPARG, SLC2A1, and TXN). (m) Results of GO enrichment analyses. (n) Results of KEGG enrichment analyses.

Then, to ascertain whether the risk score could serve as an independent prognostic factor for liver cancer patients, a multivariate Cox regression analysis was conducted. The analysis revealed that both tumor stage (HR = 1.579, 95% CI: 1.271-1.961, p < 0.001) and the risk score (HR = 1.452, 95% CI: 1.2898-1.625, p < 0.001) exhibited significant associations with overall survival time (Figure 3b). These findings demonstrate that both tumor stage and the risk score independently contribute to the prognostic prediction for liver cancer patients.

### Construction and validation of the nomogram

We constructed a nomogram model that included risk scores, age, gender, grade, and stage. The nomogram score was calculated based on a reference scale to predict the overall 1-year, 3-year, and 5-year survival rates of liver cancer patients (Figure 3c). The calibration curve of the nomogram showed good consistency between the predicted and observed results (Figure 3d).

### Analysis of model genetic differences

In gene expression analysis, the relationship between gene expression levels of these five genes and risk score was evaluated in all patient sets, the testing set, and the training set (Figure 3e-g). Analysis in each set showed that gene expression levels were correlated with risk score grouping. ADH4 had higher expression in the low-risk group compared to the high-risk group, while DLAT, PPARG, SLC2A1, and TXN had higher expression in the high-risk group compared to the low-risk group. In single gene expression analysis, there were significant differences in the expression levels of the five genes obtained from the model in both normal samples and tumor samples (Figure 3h-l). The expression level of ADH4 in tumor samples was significantly lower than that in normal samples, while the expression levels of DLAT, PPARG, SLC2A1, and TXN in tumor samples were significantly higher than those in normal samples.

### Functional enrichment analysis

We performed GO and KEGG pathway analysis on the genes with expression differences between the two risk groups. In the BP (Biological Process) category, the following were significantly enriched: mitotic sister chromatid segregation (GO:0000070; GO:0033046; GO:0033046; GO:2000816) and down-regulation of chromosome segregation (GO:0051985; GO:1905819). In the CC (Cellular Component) category, the differential genes were mainly related to the apical part of the cell (GO:0045177), the condensed chromosome, centromeric region (GO:0000779), the chromosome, centromeric region (GO:0000775), the basolateral plasma membrane (GO:0016323), the basal plasma membrane (GO:0009925), and the basal part of the cell (GO:0054178). In the MF (molecular functions) category, functions related to in vivo oxidative reduction reactions were widely enriched, such as oxidoreductase activity (GO:0016712), arachidonic acid monooxygenase activity (GO:0008391), arachidonic acid epoxygenase activity (GO:0008392), steroid hydroxylase activity (GO:0008395), heme binding (GO:0020037), and aromatase activity (GO:0070330) (Figure 3m). KEGG pathway analysis showed that the differentially expressed genes were mainly enriched in the Cell cycle (hsa04110), Lipid and atherosclerosis (hsa05417), Rheumatoid arthritis (hsa05323), Carbon metabolism (hsa01200), and other metabolic pathways (Figure 3n). In summary, the results of the enrichment analysis indicate that these 5 genes are related to cellular proliferation and immune-related metabolic pathways.

### Survival analysis in different clinical traits

The results of the survival analysis of patients in different clinical traits according to risk scores showed that the overall survival time of low-risk patients in all clinical traits was significantly higher than that of high-risk patients in the same group. This demonstrates that the scoring model is effective in predicting survival in different clinical traits (Figure 4a-h).

**Figure 4.**
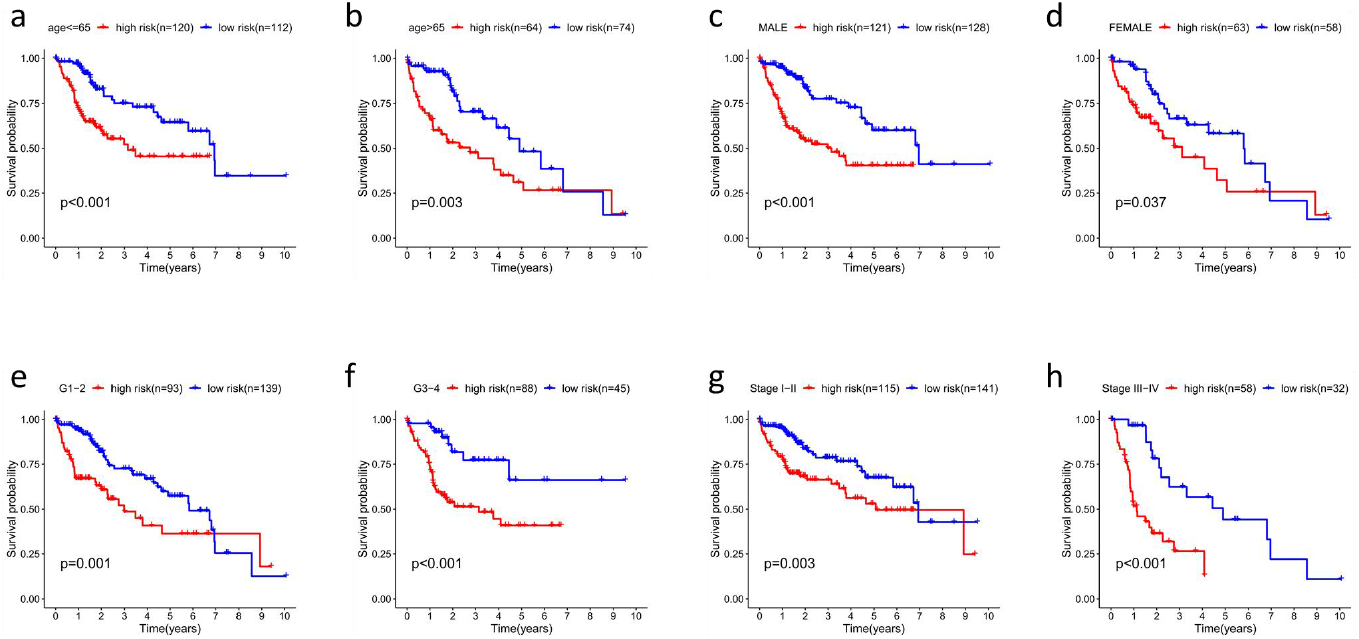
(a-h) The results of survival analysis in different clinical traits.

### Immune-related functional analysis

Immune cell infiltration is an important component of the tumor microenvironment and is closely related to tumor behavior and prognosis. The ssGSEA analysis revealed that higher risk scores were associated with significant elevation in the expression of certain immune cells (Figure 5a), including aDCs, Macrophages, Th1 cells, Th2 cells, and Tregs. However, in higher risk scores there was a lower expression of mast cells. Additionally, in immune function analysis, higher risk scores were found to be associated with increased expression of certain immune functions (Figure 5b, c), including APC_co_stimulation, CCR, Checkpoint, HLA, MHC class I, and Parainflammation. However, Type II IFN Response was suppressed in the high-risk group. Furthermore, correlations between immune-related genes and risk scores were analyzed, and the results (Figure 5d) showed that risk scores had a positive correlation with many immune-related genes, including ATIC, OLA1, CTLA4, PDCD1, CD274, IDO1, among others.

**Figure 5.**
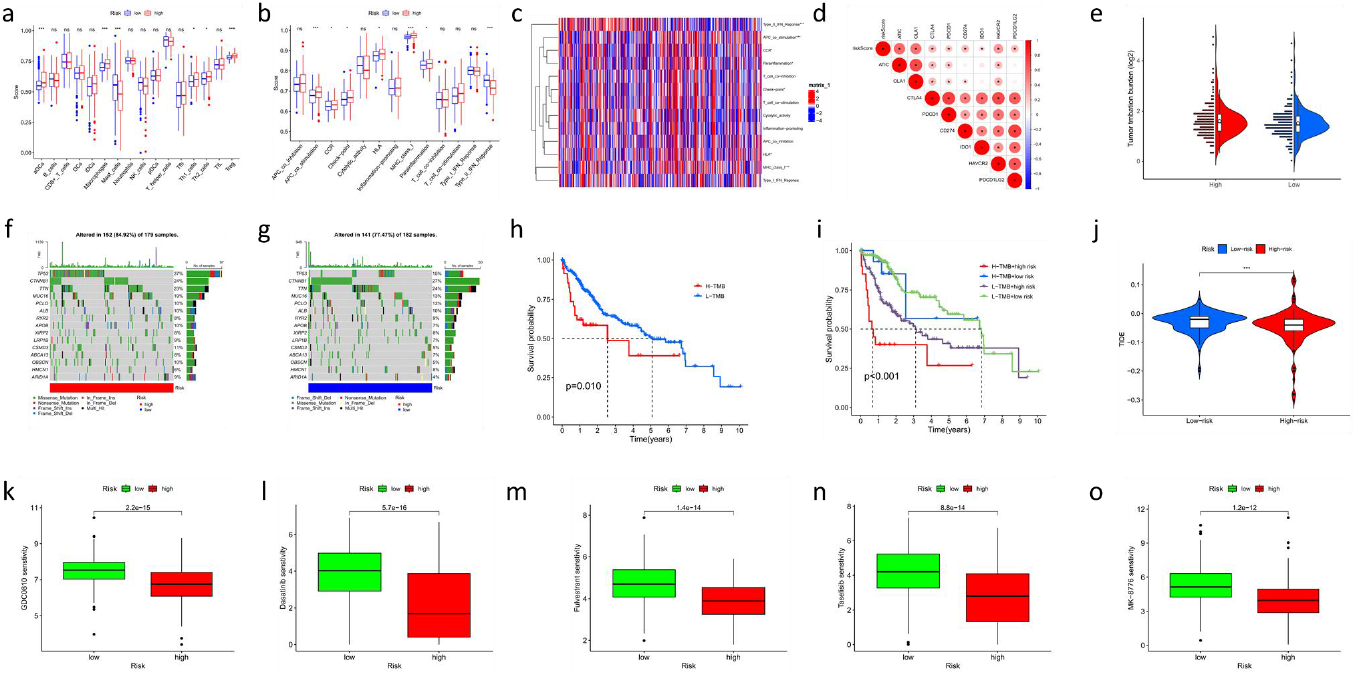
(a) The ssGSEA analysis about the scores and the expression of certain immune cells. (b) The boxplot about the scores and the immune function. (c) The heatmap analysis about the scores and the immune function. (d) The correlation between immmune-related genes and risk scores. (e) The TMB scores about the low-risk and high-risk. (f) The oncoplots of the mutation genes in liver cancer patients for the high-risk. (g) The oncoplots of the mutation genes in liver cancer patients for the low-risk. (h) The survival probibability between the high TMB groups and the low TMB groups. (i) The survival probibability with the high-low risk scores and high-low TMB. (j) The TIDE scores between low-risk and high-risk groups. (k-o) The top 5 drugs sensitity difference between low-risk and high-risk groups.

### Analysis of tumor mutation burden

Cumulative evidence suggests that Tumor Mutational Burden (TMB) is a predictive biomarker of response to immune checkpoint inhibitors (ICIs) therapy. Overall, the TMB in the high-risk group was slightly higher than that in the low-risk group, indicating a higher TMB and more gene mutations in the high-risk population (Figure 5e).

The Figure 5 f-g showed that the frequency of mutations was higher in the high-risk group than in the low-risk group (84.92% vs. 77.47%), and the mutation rate of TP53 was significantly higher in the high-risk group than in the low-risk group (37% vs. 15%). Patients in the high TMB group had shorter overall survival time than those in the low TMB group (Figure 5h). The survival rate (Figure 5i) was best for the low TMB and low-risk group and worst for the high TMB and high-risk group. When combining the risk score and TMB to evaluate the prognosis of liver cancer patients, it was found that the overall survival rate was better in the low TMB and low-risk group than in the high TMB and high-risk group. Interestingly, tumor immune evasion is lower in the high-risk group than in the low-risk group (Figure 5j), also suggesting the potential for treating liver cancer in the high-risk group through immune cell therapy.

### Drug sensitivity analysis

Five drugs were selected for sensitivity difference analysis by gene expression-based target drug prediction (Figure 5k-o). These drugs had p<0.001 with the highest expression differences between the high-risk and low-risk groups. The drugs selected were GDC0810, dasatinib, fulvestrant, taselisib and MK-8776. GDC-0810 is a nonsteroidal combination selective estrogen receptor modulator (SERM) and selective estrogen receptor degrader (SERD) (Wardell et al., 2015) for the treatment of breast cancer by blocking the HER2 signaling pathway (Cardoso, 2001; Joseph et al. 2016; Badia et al. 2022). Dasatinib is a tyrosine kinase inhibitor that inhibits BCR-ABL and SRC kinases to prevent abnormal proliferation and growth of leukemic cells (Lindauer and Hochhaus, 2018). Fulvestrant is a selective estrogen receptor degrader (SERD) that inhibits the estrogen receptor (ER) and has a higher binding affinity to the ER compared to tamoxifen (a SERM commonly used in breast cancer treatment) (Lai and Crews, 2017). Taselisib is a selective inhibitor of PI3K α that blocks downstream signaling cascade responses and induces cell death in cancer cells (Zumsteg et al., 2015). Mk-8776 is a selective inhibitor of checkpoint kinase 1 (CHK1) that sensitizes cancer cells to chemotherapy-induced DNA damage and promotes cell death (Bridges et al., 2016). In summary, gene expression-based target drug prediction identified five drugs for differential sensitivity analysis. These drugs, including GDC0810, dasatinib, fulvestrant, taselisib, and MK-8776, showed promising results in preclinical models of cancer and may improve treatment outcomes and reduce the incidence of drug resistance. Further studies are needed to optimize their dosing and to identify biomarkers that predict treatment response.

## Discussion

Liver cancer is a global health concern, with increasing incidence rates and significant mortality worldwide. In our study, we investigated the potential of NAD^+^ metabolism-related genes as novel therapeutic targets for liver cancer. We identified key genes among NAD^+^ metabolism-related genes and explored their relationship with cancer staging and prognosis based on gene expression levels.

Our study successfully constructed a risk score model using five NAD^+^ metabolism-related genes, providing a quantitative measurement of cancer risk and survival status. The risk score model demonstrated significant predictive potential for survival outcomes, outperforming traditional clinical features such as age, gender, grade, and stage. These findings highlight the clinical value and significance of our predictive model in liver cancer prognosis prediction.

The prognostic significance of the five selected NAD^+^ metabolism-related genes was further confirmed through various analyses. We observed distinct expression patterns of these genes between high-risk and low-risk patients, both in overall patient sets, testing set, and training set. Additionally, the expression levels of these genes showed significant differences between normal samples and tumor samples, further indicating their relevance in liver cancer.

Functional enrichment analysis revealed that these five genes are involved in cellular proliferation and immune-related metabolic pathways. This suggests their potential role in tumor development and immune evasion mechanisms. Immune-related functional analysis demonstrated associations between the risk score and immune cell infiltration, indicating potential implications for immunotherapy strategies in liver cancer treatment.

Furthermore, we explored the relationship between risk score and tumor mutation burden (TMB). The high-risk group exhibited a higher TMB and more gene mutations, potentially influencing the response to immune checkpoint inhibitors (ICIs) therapy. Drug sensitivity analysis identified five drugs that showed differential sensitivity between high-risk and low-risk groups. These drugs, including GDC0810, dasatinib, fulvestrant, taselisib, and MK-8776, hold promise for improving treatment outcomes and reducing drug resistance.

In summary, our study highlights the importance of NAD^+^ metabolism-related genes in liver cancer prognosis prediction. The risk score model constructed based on these genes demonstrates significant predictive potential and clinical value. The identification of key genes and their associations with cancer risk, survival outcomes, immune-related functions, and drug sensitivity provide valuable insights for developing personalized therapeutic strategies for liver cancer patients. Further research is warranted to validate and refine these findings, ultimately improving the management and outcomes of liver cancer.

## Conclusion

In conclusion, our study demonstrates the potential of NAD^+^ metabolism-related genes as prognostic markers and therapeutic targets in liver cancer. We successfully constructed a risk score model using five key genes, which exhibited significant predictive power for survival outcomes in liver cancer patients. These genes play crucial roles in tumor development, immune evasion, and metabolic pathways. Additionally, the risk score showed associations with tumor mutation burden and differential drug sensitivity, indicating its potential for guiding personalized treatment approaches. Our findings provide valuable insights into the molecular mechanisms underlying liver cancer and pave the way for the development of targeted therapies. Further research and validation are needed to fully exploit the clinical implications of NAD^+^ metabolism-related genes in liver cancer management.

## Notes

### Competing Interest Statement

The authors have declared no competing interest.

